# PEAR: a flexible fluorescent reporter for the identification and enrichment of successfully prime edited cells

**DOI:** 10.1101/2021.04.26.441486

**Authors:** Dorottya A. Simon, András Tálas, Péter I. Kulcsár, Ervin Welker

**Affiliations:** Institute of Enzymology, Research Centre for Natural Sciences, Budapest, Hungary; ProteoScientia Ltd., Hungary; School of Ph.D. Studies, Semmelweis University, Budapest, Hungary; Biospiral-2006 Ltd., Szeged, Hungary; Institute of Biochemistry, Biological Research Centre, Szeged, Hungary

## Abstract

Prime editing is a recently developed gene engineering tool that allows the introduction of short insertions, deletions or substitutions into the genome. However, the efficiency of prime editing, generally reaching around 10-30% editing, has not resembled its versatility. Here, Prime Editor Activity Reporter (PEAR), a sensitive fluorescent tool is introduced for the identification of single cells with prime editing activity. Possessing no background fluorescence, PEAR specifically reports on prime editing events in individual cells. By design, it ensures unrestricted flexibility for sequence variations in the full length of the target sequence. The application of PEAR as an enrichment marker of prime editing can increase the edited population by up to 70% and alleviate the burden of the otherwise time and labour consuming process of cloning of the correctly edited cells, therefore considerably improving the applicability of prime editing in fundamental research and biotechnological uses.

## Introduction

Discovery of CRISPR systems in bacteria and archaea has not only drastically increased the spectrum of organisms that we can genetically modify but it has also equipped us with a simple tool for the introduction of a variety of different types of modifications into the genome ^[1–3]^. Many of the CRISPR-based approaches rely on generating double strand breaks that is usually accompanied by genome-wide off-target editing ^[4–7]^. Recently, more precise CRISPR tools; base ^[8,9]^ and prime ^[10]^ editors have been developed, that can introduce modifications into the DNA in a base pair resolution without the requirement of donor DNA templates or the introduction of double-strand DNA breaks. While current base editor (BE) variants provide efficient editing, they are restricted to certain substitution mutations ^[8,9,11,12]^. In contrast, prime editors (PEs) can introduce all types of substitutions and/or precise indels, however, their efficiency lags behind of BEs’. PEs consist of a nickase version of *Streptococcus pyogenes* Cas9 (SpCas9) fused to a reverse transcriptase enzyme ^[10]^. The fused reverse transcriptase can extend the nicked DNA strand using an RNA template that is located on the 3’ terminus of an extended, single guide RNA (sgRNA), called the prime editing sgRNA (pegRNA). By careful design of the pegRNA, modifications can be introduced downstream of the nick generated on the non-targeted DNA strand. The 3’ end of the pegRNA, that is complementary to the non-targeted strand, consists of the reverse transcriptase template (RT) and the primer binding site (PBS). The PBS is complementary to the 3’ end of the nicked non-targeted DNA strand, with which it forms a hybrid DNA-RNA helix. The RT, which contains the mutation(s) to be introduced, also comprises the protospacer adjacent motif (PAM). The optimal length of the PBS and the RT varies from target to target and requires extensive optimisation for each different target. The efficiency of prime editing can be increased by nicking the non-edited strand (prime editor 3 – PE3) at the expense of generating unwanted indels at the targeted locus ^[10]^. The effect of the nicking varies depending on its position and it requires further optimisation. In some cases, the efficiency of prime editing can be further increased by also altering the PAM sequence with the desired mutation (PE3b); this may prevent the SpCas9 from cleaving the sequence again. This method can reduce the chance of co-introducing unwanted indels with the intended mutations, thus increasing the occurrence of precise modifications ^[10]^.

The development of improved PE variants can be forecasted based on the history of base editors that are now available with highly improved features and efficiency ^[13–19]^. The aim of our study was to develop a reporter system for an easy, fluorescence-based detection of prime editing outcomes, that allows maximum flexibility for the target sequences and pegRNA designs to be tested on it. Several systems have been developed for reporting base editing that use BEs for the generation or alteration of a start or stop codon^[20,21]^, or to rescue a disruptive amino acid and subsequently recover functions of antibiotic resistance genes or fluorescence proteins ^[22]^. Alternatively, a non-synonymous mutation in the chromophore of a fluorescent protein that induces fluorescence spectral change has also been explored as a method to monitor base editing activity ^[23,24]^. Reporter systems have also been demonstrated for prime editing, but they are either restricted to few target sequences ^[25,26]^ and/or showed a rather low signal ^[20]^.

We also wanted to find out whether prime edited mammalian cells could be identified and enriched using a plasmid-based surrogate marker for chromosomal DNA modifications, as demonstrated before with Cas nucleases ^[27,28]^ and base editors ^[20,23,29,30]^.

## Results

We aimed to develop a reporter system, that possesses several key features. It should be a transient plasmid-based system that is not restricted to one or a few cell lines nor does it require extensive work, such as the generation of cell lines. It needs to be based on a gain-of-function fluorescent signal with minimal background to be capable of detecting the signal in the timeframe of a transient system. The sequence requirements of efficient prime editing are not yet fully understood ^[10,31]^; thus, it is crucial for the widespread application of a reporter system, that the sequence of the target and the flanking nucleotides of the position to be edited can be freely interchanged. We have recently developed a base editor activity reporter (BEAR), which meets these criteria ^[30]^, and it has the potential to be converted into a suitable tool for prime editing. BEAR is based on a split GFP protein separated by the last intron of the mouse *Vim* gene. The sequence of the functional 5’ splice site is altered in such way, that splicing and therefore the GFP fluorescence is disrupted, however, they can be restored by applying base editors (**Figure S1a, b**). In this system the G-GT-RAGT sequence, which ensures optimal splicing at the 5’ splice site, can be altered by one nucleotide without a significant decrease in the splicing efficiency. A systematic investigation of the sequence requirements for efficient splicing has revealed, that the activity of several inactive splice variants could be recovered by restoring just one nucleotide in its sequence, using either adenine base editors (ABEs) or in other cases, cytosine base editors (CBEs). Using the information acquired during the development of the BEAR system ^[30]^ and exploiting the ability of prime editing to alter more than one nucleotide simultaneously, we have designed a plasmid that contains an inactive splice site, which can be activated by the action of a prime editor harbouring an appropriately designed pegRNA (**Figure 1a,b**). Cells containing plasmids with an active splice site sequence will then be able to efficiently express GFP, which can be quantified by flow cytometry. Hence, we name assays that exploit this design as Prime Editor Activity Reporters (PEARs).

**Figure 1 –.**
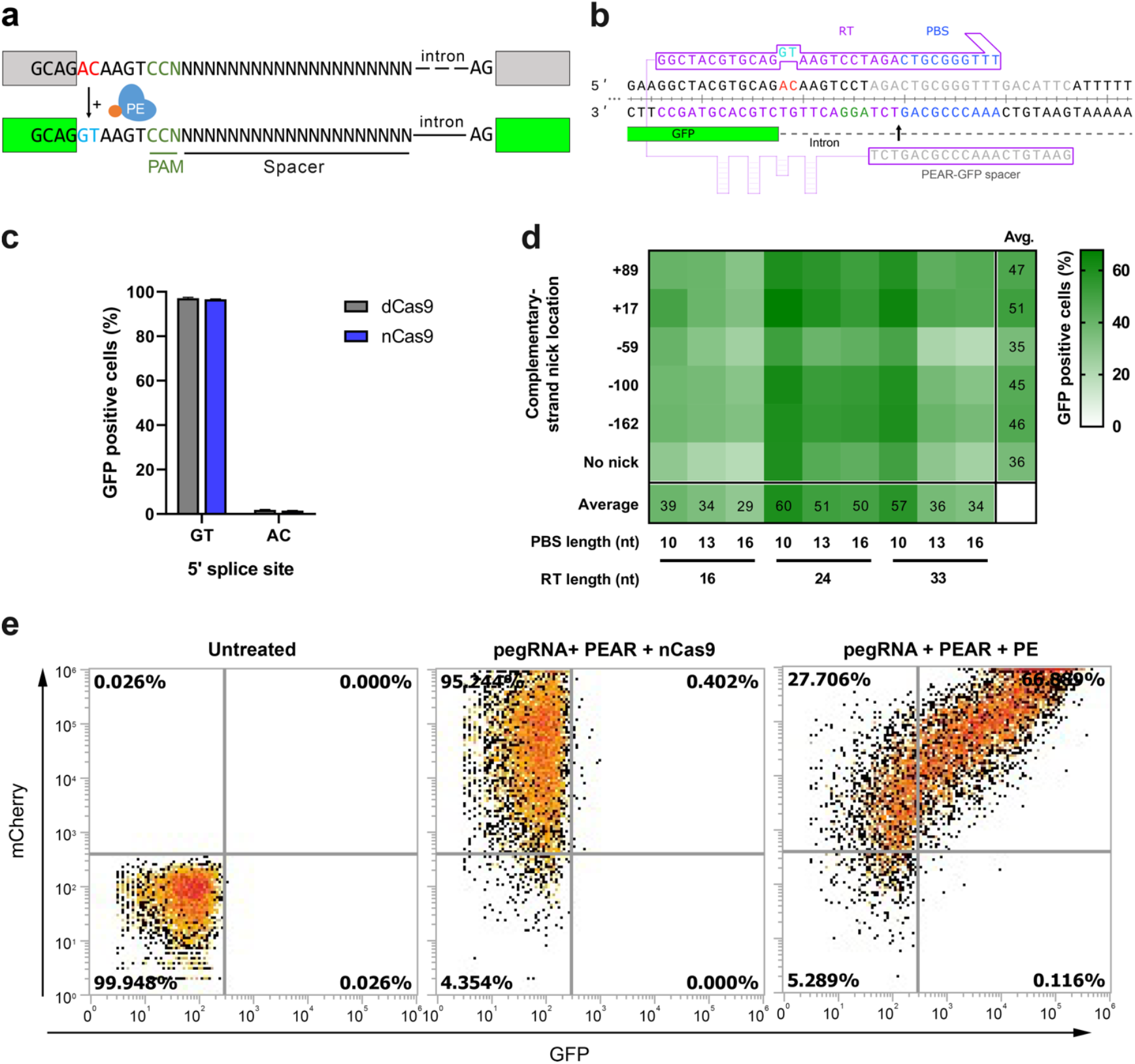
Principle of the Prime Editor Activity Reporter (PEAR) assay. (**a**) Schematic of the Prime Editor Activity Reporter (PEAR). The mechanism of PEAR is based on the same concept as BEAR, and it contains the same inactive splice site, shown in panel (a). PE can revert the ‘G-AC**-**AAGT’ sequence to the canonical ‘G-GT-AAGT’ splice site. Unlike BEAR, here prime editing occurs on the antisense strand of the DNA, hence, this method enables us to position the spacer sequence within the intron. Here, the entire length of the spacer is freely adjustable (shown as ‘N’-s). The altered bases of the splice site are shown in red, and the edited bases are shown in blue. The PAM sequence is dark green, nCas9 is blue and the fused reverse transcriptase is orange. (**b**) Detailed view of the 5’ splice site and the surrounding sequences of the PEAR-GFP plasmid. The 3’ end of the first exon of GFP is shown in green, the intron is shown as a dashed line. The spacer sequence in the pegRNA and the target sequence in the DNA are shown in grey, the PAM is green, and the inactive splice site is red. The RT (purple) and the PBS (blue) sequences in the pegRNA and the targeted sequences are also coloured. The Cas9 nick site is indicated by a black arrow. (**c**) PEAR-GFP plasmids with an active or inactive 5’ splice site. Flow cytometry measurements of GFP positive HEK293T cells co-transfected with a PEAR-GFP reporter plasmid harboring a pre-edited active (GT) or an inactive (AC) 5’ splice site and nuclease (nCas9, blue) or dead Cas9 (dCas9, grey) constructs as indicated in the figure. (**d**) Optimisation of PBS, RT and complementary DNA strand nicking on the PEAR-GFP plasmid. The heatmap shows the average percentage of GFP positive cells of three replicates of transfections with the PEAR-GFP plasmid and prime editor, using different pegRNAs and secondary nick locations. The position of the second nick is given in relation to the first nick. Positive values indicate 3’, negative values indicate 5’ direction on the targeted DNA. When no second nick was introduced it is indicated as “no nick”. (**e**) Flow cytometry measurements of control and prime edited HEK293T cells using the PEAR-GFP plasmid. Density dot plots of the non-transfected (left), nickase Cas9 transfected (middle) and PE transfected (right) cells. The pegRNA plasmid also expresses mCherry for monitoring transfection. In the plots in the middle and on the right, a pegRNA with an RT length of 24 and PBS length of 10 was used, and the second nick was introduced in position +17.

The editing windows of base editors exploited for the BEAR system are located PAM distal, 5’ of the nick (**Figure S1a,b**). PEs on the other hand can introduce mutations 3’ of the nick, a region that contains the PAM sequence (**Figure 1a,b**). To ensure maximum flexibility for the SpCas9 target, we kept the target sequence in the intron by moving the PAM to the complementary strand. As a result, the target sequence can be freely adjusted in its entire length (**Figure 1a,b**). Thus, while BEAR could be used with a few millions of different target sequences, PEAR offers unrestricted sequence variations for the SpCas9 target. Additionally, PEAR preserves the flexibility of BEAR to interchange the targeted and the flanking nucleotides ^[30]^. No other fluorescence-based reporter systems have demonstrated such degree of flexibility before ^[20,25,26]^.

HEK293T cells were transfected with either an inactive or a nickase SpCas9 (dSpCas9 or nSpCas9, respectively) along with plasmids containing either the inactive splice site sequence selected to be edited (PEAR-GFP), or a functioning one, which would result from the action of a PE. The latter plasmid harbouring the functional 5’ splice site is hereafter referred to as pre-edited. Cells showed high fluorescence when dSpCas9 (as a negative control) was co-transfected with the pre-edited plasmid while the PEAR-GFP with disrupted 5’ splice site resulted in no expression of the fluorescent protein (**Figure 1c)**. When the two plasmids were co-transfected with the H840A nickase Cas9 (a component of the prime editor) it did not alter the fluorescence of cells compared to the negative control (**Figure 1c)**. These results are in line with former observations that indels cannot, only substitution mutations in the inactive 5’ splice site can restore fluorescence ^[30]^ and suggest that this system will likely be suitable to report specifically on the action of prime editors.

To see whether PE can indeed recover fluorescence using the PEAR-GFP plasmid, several pegRNAs combining various PBS- and RT-lengths and complementary nicking positions were tested. The heatmap in **Figure 1d** shows that the PEAR-GFP plasmid can in fact be efficiently edited under several conditions. The editing shown in **Figure 1e** applying ten nucleotide-long PBS (PBS-10) and the 24 nucleotide-long RT region (RT-24) gave the highest efficiency among the tested combinations. The sixth row of the heatmap in **Figure 1d** shows the editing efficiency when the complementary strand is not nicked. In line with the expectations, these values are generally smaller, than when the complementary strand is nicked ^[10]^. The best complementary strand nicking site is position +17 (in relation to the first nick) in the case of most PBS and RT length combinations. Interestingly, in all conditions PBS-10 gives greater values than PBS-13 and PBS-16 when combined with any of the RTs. Comparing the RTs, the efficiency of the editing is in the order of RT-24>RT-33>RT-16 in most conditions (16 out of 18). These results are consistent with the concept that the effect of the length of PBS and RT on editing efficiency is primarily independent from one another ^[32]^. It is also in line with the current optimisation practice, where first, the length of the PBS is optimised using a given RT length, and then the length of the RT is optimised using the PBS selected in the first step. Thus, these experiments support the hypothesis, that PE efficiency is governed by the same factors when using a PEAR plasmid, as demonstrated earlier on chromosomal targets ^[10,31]^.

In order to validate this idea, we compared prime editing efficiency on PEAR plasmids with two HEK293T cell lines (that were generated in ref. ^[30]^); now we named them as HEK-PEAR-GFP and HEK-PEAR-mScarlet, that contain a chromosomal copy of a PEAR plasmid (PEAR-GFP-2) harbouring either an interrupted GFP or an mScarlet sequence, respectively. To facilitate the comparison, we exploited the corresponding plasmids (PEAR-GFP-2 and PEAR-mScarlet plasmids, respectively) harbouring the same sequence surrounding the inactive splice site in the cell lines. Since the sequences to be edited in the two cell lines were designed to be used with base editing that works 5’, upstream of the PAM sequence, we needed to identify another PAM site that would allow the application of prime editing. **Figure 2** and Figure S2 shows the layout of PEAR-GFP-2 (**Figure 2a)** and PEAR-mScarlet (**Figure 2d, Figure S2a)** with the target and secondary nick positions clearly indicated. The two nucleotides of the inactive splice site to be corrected are in positions +16-17 in relation to the new PAM in HEK-PEAR-GFP. In the HEK-PEAR-mScarlet cell line they are in positions +26-27 or +33-34 in relation to the two PAMs.

**Figure 2 –.**
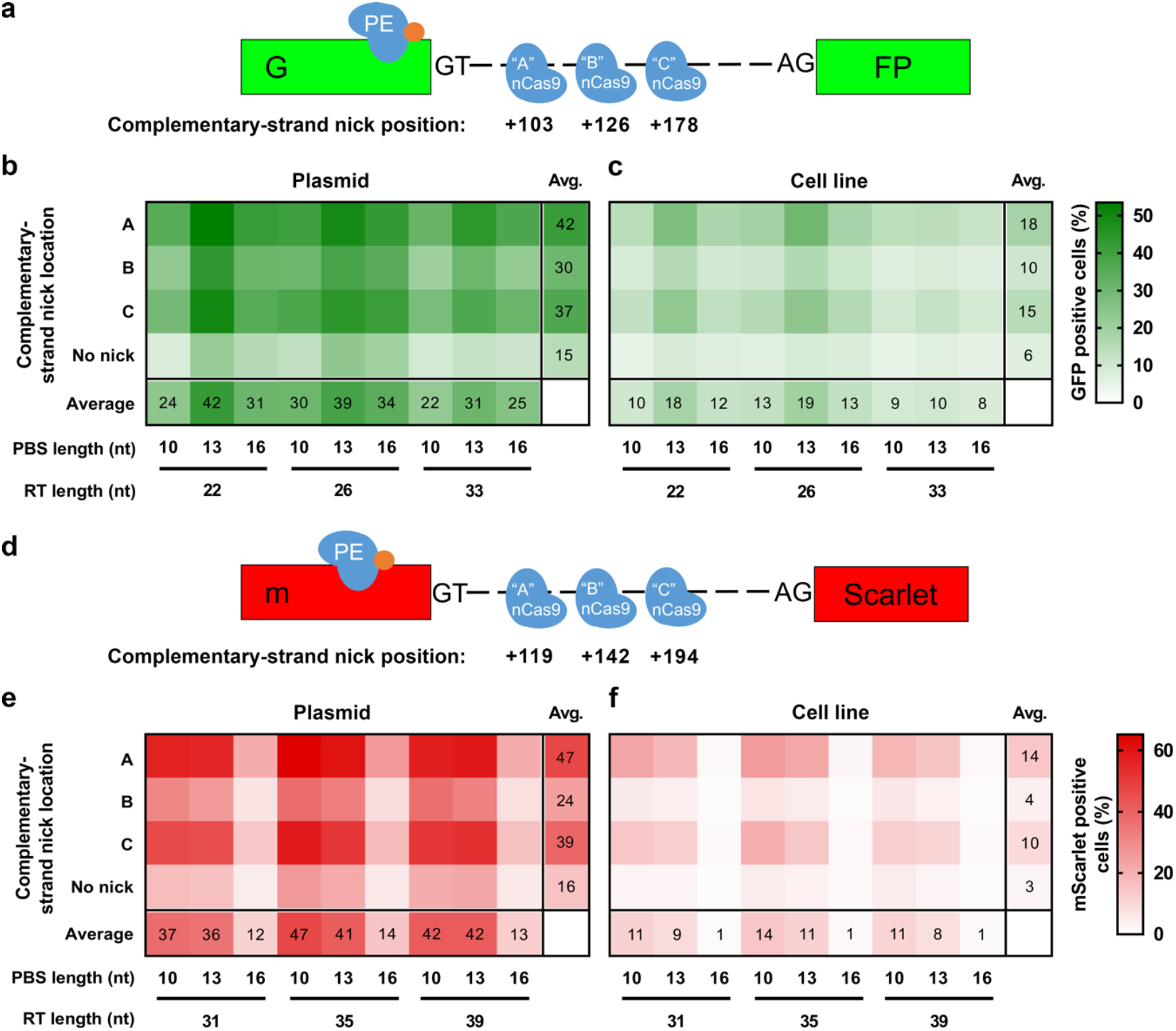
Prime editing with PEAR in a genomic context. Schematic diagram of the PEAR-GFP-2 (**a**) and the PEAR-mScarlet (**d**) coding sequence is shown alongside the cell lines in panels (**c** and **f**), which contain the corresponding plasmid. The prime editor corrects the inactive splice site sequence and restores GFP or mScarlet fluorescence, respectively. The second nick positions that enhance prime editing are the same in (**a**) and (**d**), and are indicated as A, B or C. The heatmaps show the average percentage of GFP (**b** and **c**) and mScarlet (**e** and **f**) positive cells derived from three replicates, where the prime editor was transfected into cells alongside different pegRNAs and sgRNAs with different complementary strand nick locations. In panel (**b**) PEAR-GFP-2 and in panel (**e**) PEAR-mScarlet plasmid is co-transfected. In panel (**c**) HEK-PEAR-GFP and in panel (**f**) HEK-PEAR-mScarlet is the transfected cell line. On the left side of the heatmaps the positions of the second nicks are indicated with A, B and C letters, corresponding to those in panel (**a**) and (**d**). When no second nick was introduced, it is indicated as “no nick”.

In case of the GFP expressing cells, we tested 36 different conditions for both the cell line and the PEAR-GFP plasmid: all possible PBS – RT – second nick site combinations. The heatmap in **Figure 2b** shows that PBS-13 and RT-22 or RT-26 in combination with the second nick site at position 103 results in maximum efficiency for the PEAR-GFP-2 plasmid. In lack of the second nick, the efficiency of the editing is considerably lower. Comparing these results with the efficiency of prime editing in the cell line (**Figure 2c)**, the most ideal conditions are the same. The overall pattern of the two heatmaps is indisputably similar, exhibiting strong correlation (r=0.89). In **Figure 2c a** somewhat lower efficiency is apparent in general compared to that apparent with the plasmid in **Figure 2b**. This difference is likely due to the much higher copies of the plasmids that are present in the cell during the experiments in **Figure 2b**.

In case of the mScarlet expressing cells, first, we compared the efficiencies of prime editing with target 1 (**Figure S2b)** and target 2 (**Figure S2c)**, by exploring 36 different combinations of PBS, RT and second nick site for each target employing only the corresponding PEAR-mScarlet plasmid. Target 1 gave higher prime editing efficiencies and with both targets PBS 10 and 13 was optimal, but PBS 16 had a negative effect on prime editing efficiency.

With target 1 the top working PBS is 10 nucleotides with both the fluorescent cell line (**Figure 2f)** and the PEAR-mScarlet plasmid (**Figure 2e)**, the differences between the various combinations are more apparent with the cell line. The two heatmaps exhibit a remarkably similar pattern and a strong correlation can be seen between the editing efficiency using the cell line and the PEAR plasmid (r=0.93). Similar results (r=0.92) can be achieved with target 2 when comparing plasmid (**Figure S2d)** and genomic (**Figure S2e)** results. Collectively, these results strongly imply that the main features of prime editing and factors affecting its efficiency are accurately reflected within our system.

Post-transfection selections of edited cells by FACS or antibiotics have proven to be effective methods for the enrichment of genetically modified cells within a population, provided that the Cas9 plasmid co-expresses a protein that is fluorescent, or it bears antibiotic resistance ^[20,24,27–30]^. Experiments also showed that selecting for cells with a surrogate marker subjected to the same type of genetic modification (knockout, HDR or base editing), results in higher enrichment of successfully edited cells, than when solely selecting for transfection markers ^[20,23,27–30]^ suggesting that some cells allow for higher editing efficiency. This phenomenon could be due to the nuclease being able to exhibit higher activity, or the activity of the DNA repair system could be considerably more engaged in these cells. We proposed that the level of enrichment, which can be achieved using the transfection marker alone, can be exceeded by enriching cells in which a marker protein in a plasmid is simultaneously prime edited. The PEAR-GFP plasmid was used in these experiments as a surrogate marker with previously optimised RT and PBS conditions (**Figure 1d)**. To test this principle, first, we used the PEAR-mScarlet cell line and measured the ratio of the mScarlet positive cells in (1) all measured cells, (2) within cells that either express the blue fluorescent protein (BFP) for transfection enrichment or (3) express both BFP and GFP for PEAR enrichment. For this purpose, we chose two efficient previously tested PEAR-GFP targeting pegRNAs (**Figure 1d)**. **Figure 3a** shows that transfection marker enrichments resulted in a significant increase in the ratio of edited cells to non-edited cells. Gating for the BFP-expressing cells increased the percentage of the edited population from 18% to 24% with both pegRNAs tested. PEAR-enrichment resulted in a considerable increase compared to the transfection enrichments in case of both pegRNAs (57 and 76% mScarlet expressing cells, respectively). These results demonstrate that prime edited cells can be substantially enriched when using PEAR plasmid as a surrogate marker.

**Figure 3 –.**
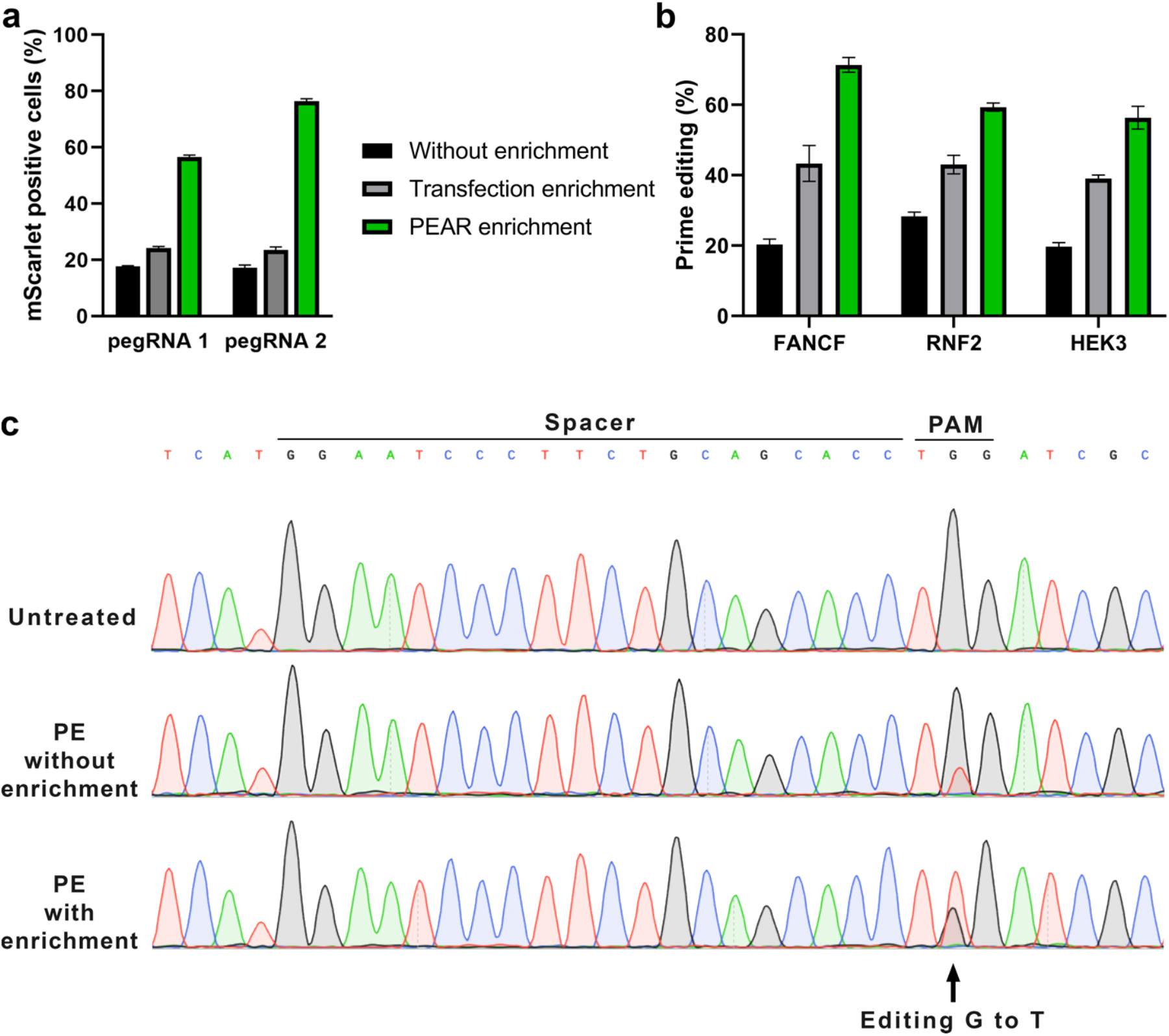
PEAR enriches for prime edited cells. (**a**) Flow cytometry measurements of prime edited (i.e., mScarlet positive) cells in the HEK-PEAR-mScarlet cell line. The PEAR-GFP plasmid, prime editor and two pegRNAs targeting either the plasmid or the genomic PEAR sequences were co-transfected with sgRNAs (for complementary strand nicking). Two efficient pegRNAs were selected from a previous experiment (**Figure 1d**) to edit the PEAR-GFP plasmid (pegRNA 1 had an RT of 24 and a PBS of 10, pegRNA 2 had an RT of 33 and a PBS of 10). The mScarlet positive cell count was determined from cells that were gated for all live single cells (black bars), for BFP positive cells (transfection enrichment, gray bars), and for both BFP and GFP positive cells (PEAR enrichment, green bars). (**b**) FACS enrichment of cells bearing prime edited targets. EditR and Sanger sequencing were used to assess genome editing efficiency. Percentages of the desired base editing outcome is shown. The PEAR-GFP plasmid and endogenous genomic targets (FANCF, RNF2 and HEK3) were simultaneously edited by PE. Edited cells were sorted to 3 fractions: all cells (without enrichment, black), mCherry positive cells (transfection enrichment, grey), and both mCherry and GFP positive cells (PEAR enrichment – green). Columns represent means +/−SD of three replicates. (**c**) Chromatograms of Sanger sequencing of the FANCF genomic site are shown with or without prime editing and/or PEAR enrichment. The expected G to T modification’s position 5 nucleotides from the nick is indicated by an arrow.

To test the applicability of PEAR on enriching the prime edited population in endogenous targets, we chose three genomic targets (FANCF, RNF2, HEK3) for which the optimal pegRNAs are already known ^[10]^, and using these targets, editing is expected to be readily detectable with Sanger sequencing and EditR, even without enrichment (**Figure 3b, c**). Here, mCherry was used as the indicator of transfection efficiency, and the PEAR-GFP plasmid was employed here too as a surrogate marker. On the third day from transfection, cells were sorted into three fractions: (1) all single, living cells; (2) mCherry positive cells, and (3) mCherry and GFP positive cells. After genomic DNA purification from the populations and PCR amplification of the targeted amplicon, the editing percentage was determined using EditR: G to T, C to A and T to A editing were detected at the FANCF, RNF2 and HEK3 sites, respectively. The editing rates have increased using transfection enrichment (from 20% to 43%, 28% to 43% and 19% to 39%, respectively), however, they have all showed a greater increase when the PEAR enrichment was used (to 71%, 59% and 56%, respectively). These results demonstrate that PEAR can substantially enrich for prime edited cells in transfected populations, reaching about 70% prime-edited cells.

## Discussion

Prime editing provides a great flexibility for genomic modifications without having to supply a donor DNA or risking a double strand DNA break ^[10]^. This technology could revolutionise the rapidly expanding universe of genome editing. The prime editing technology has readily been translated from mammalian models ^[10,26,32–34]^ to plants ^[25,35–37]^ and *Drosophila* ^[38]^. Despite its great potential, we have found surprisingly few studies that had demonstrated the practical use of prime editing in mammalian cells ^[26,34]^ which is a vast underestimation of the true flexibility, that this method promises. This might be due to the extensive optimisation it requires and the relatively low editing efficiency that can be achieved at a subset of targets. In this aspect, any approach that can substantially increase the editing efficiency is critical to aid its general use. Here, we have developed PEAR, a sensitive fluorescent assay for the monitoring of prime editing activity in mammalian cells, and we have also shown that prime editing occurs at much higher frequencies in those individual cells, where the PEAR plasmid is co-edited. In fact, the enrichment with a transfection marker doubled, while PEAR enrichments tripled the number of edited cells in our experiments (**Figure 3a-c**). Thus, the application of PEAR as a surrogate marker could substantially contribute to the more widespread use of prime editing, by considerably increasing the number of potential targets, which may now be edited with appropriate efficiency.

PEAR is also a versatile tool that can be utilised for the characterisation of prime editors. We examined more than 250 conditions combining different sgRNAs, PBS- and RT-lengths and nick sites, and our results confirmed that the efficiency of prime editing to modify PEAR plasmids is governed by the same factors as prime editing on chromosomal targets (**Figure 2)**. The tolerance of the 5’ splice site for substitutions ^[30]^ makes the sequences of target region of the PEAR plasmid easily adjustable. This versatility allows the user to examine the effect of sequence features and other factors affecting the efficiency of prime editing in a systematic manner that cannot be explored otherwise using genomic targets. Our approach beyond its general use may also assist the development of more efficient prime editors or variants with enhanced characteristics in the future.

Owning to the relatively low efficiency of prime editing with many potential target sequences, PEAR is expected to become a widespread tool to be used in a wide variety of biomedical and biotechnological applications. PEAR may be an especially useful tool to be used with hard-to-transfect cells or where the efficiency of PEs is particularly compromised.

## Materials and Methods

### Materials

Restriction enzymes, T4 ligase, Dulbecco’s modified Eagle medium (DMEM), fetal bovine serum, Turbofect and penicillin/streptomycin were purchased from Thermo Fischer Scientific Inc. DNA oligonucleotides and the GenElute HP Plasmid Miniprep and Midiprep kit used for plasmid purifications were acquired from Sigma-Aldrich. Q5 High-Fidelity DNA Polymerase, NEB5-alpha competent cells, HiFi Assembly Master Mix were purchased from New England Biolabs Inc.

### Plasmid construction

The PEAR-GFP plasmid (pDAS12125_PEAR-GFP) was constructed by Esp3I digestion of the pAT9624-BEAR-cloning plasmid ^[30]^, followed by one-pot cloning of the linker oligonucleotides *(12125-L1 and –L2)*. The reaction included 2 units of Esp3I enzyme, 1.5 units of T4 DNA ligase, 1 mM DTT, 500 μM ATP, 50 ng vector and 5-5 μM of target-coding oligonucleotides. Components were mixed in with 1x Tango buffer, and the mixture was incubated at 37°C for 30 minutes before being transformed into NEB5-alpha competent cells. The pre-edited PEAR-GFP plasmid (pDAS12124_PEAR-GFP-preedited) was constructed by the one-pot cloning protocol above, using linker oligonucleotides *12124-L1 and –L2*. Previously used plasmids pAT9651-BEAR-GFP and pAT9752-BEAR-mScarlet ^[30]^ were renamed here, for consistent nomenclature, to PEAR-GFP-2 and PEAR-mScarlet, respectively, since these plasmids were used here for the evaluation of prime editing.

To monitor transfection efficiency, fluorescent protein (mCherry or TagBFP) expressing pegRNA cloning plasmids were constructed (pDAS12069-U6-pegRNA-mCherry, pDAS12070-U6-pegRNA-BFP). To construct both plasmids, an exchangeable cassette from a previously made plasmid was cloned into pAT9658 and pAT9679 to NdeI and EcoRI sites. Both pegRNA cloning plasmids bear a spacer cloning site (between BpiI sites) and a PBS-RT cloning site (between Esp3I sites).

For the construction of pegRNAs, a one-step pegRNA cloning protocol was described by Anzalone et al. ^[10]^, however, it resulted in frequent deletions and/or mutations in the constructs, therefore we also used alternative strategies. The most effective was a two-step cloning procedure: first, the spacer coding linkers were cloned into pDAS12069 or pDAS12070 plasmids between BpiI sites using 2 units of the BpiI enzyme, 1.5 units of T4 DNA ligase, 500 μM ATP, 1x Green buffer, 50 ng vector and 5-5 μM of each annealed oligonucleotides. In the second cloning step the PBS-RT bearing oligonucleotides were cloned into between Esp3I sites of the plasmids created in the first cloning step with the above method using 1 mM DTT and 1X Tango buffer. The RNF2 targeting pegRNA in Figure 3 has a C base in position 83.

For the second nicking of a plasmid or the genome, all sgRNA targets were cloned into pAT9658-sgRNA-mCherry or pAT9679-sgRNA-BFP plasmids between BpiI sites via one pot cloning, as described above.

For detailed plasmid (**Table S1**) and primer (**Table S2**) information see Supporting Information. The sequences of all plasmid constructs were confirmed by Sanger sequencing (Microsynth AG).

Plasmids acquired from the non-profit plasmid distribution service Addgene were the following: pCMV-PE2 (#132775, ^[10]^), pAT9624-BEAR-cloning (#162986), pAT9651-BEAR-GFP (#162989), pAT9752-BEAR-mScarlet (#162991), pAT9658-sgRNA-mCherry (#162987), pAT9679-sgRNA-BFP (#162988) -^[30]^, pX330-Flag-wtSpCas9-H840A (#80453), pX330-Flag-dSpCas9 (#92113).

The following plasmids developed in this study are available from Addgene: pDAS12125_PEAR-GFP (#00000), pDAS12124_PEAR-GFP-preedited (#00000), pDAS12069-U6-pegRNA-mCherry (#00000), pDAS12070-U6-pegRNA-BFP (#00000), pDAS12230_pegRNA-PEAR-GFP(10PBS-24RT)-mCherry (#00000), pDAS12137_sgRNA-PEAR-GFP-nick(+17)-mCherry (#00000).

### Cell culturing and transfection

HEK293T (ATCC, CRL-1573) cells were grown at 37°C in a humidified atmosphere of 5% CO_2_ in DMEM medium supplemented with 10% heat-inactivated fetal bovine serum with 100 units/mL penicillin and 100 μg/mL streptomycin.

HEK293T cells were seeded on 48-well plates a day before transfection, at a density of 5 × 10^4^ cells/well. A total of 565 ng DNA was used: 55 ng of PEAR target plasmid, 153 ng of pegRNA-mCherry (or pegRNA-BFP in the case of PEAR-mScarlet), 49 ng of sgRNA-mCherry (or sgRNA-BFP in the case of PEAR-mScarlet), and 308 ng of PE2 coding plasmid. These were mixed with 1 μl Turbofect reagent diluted in 50 μl serum-free DMEM, and the mixture was incubated for 30 minutes at RT before adding it to the cells. Each transfection was performed in three replicates. Cells were analysed by flow cytometry on day three, after the transfection.

The BEAR-GFP and BEAR-mScarlet cell lines were constructed as described earlier in ref. ^[30]^. The previously used BEAR-GFP and BEAR-mScarlet cell lines were renamed here, for consistent nomenclature, to PEAR-GFP and PEAR-mScarlet cell lines, since these were used here for the evaluation of prime editing. These stable cell lines were transfected with a total of 565 ng DNA: 170 ng of pegRNA-mCherry (or pegRNA-BFP in the case of PEAR-mScarlet), 54 ng of sgRNA-mCherry (or sgRNA-BFP in the case of PEAR-mScarlet) and 340 ng of PE2 coding plasmid, that was all mixed with 1 μl Turbofect reagent diluted in 50 μl serum-free DMEM. The mixture then was incubated for 30 minutes at RT before adding it to the cells.

In the enrichment experiments, where fluorescence activated cell sorting was used, HEK293T cells cultured on T-25 flasks were seeded a day before transfection at a density of 1.3 × 10^6^ cells/flask. 7924 ng total DNA: 1754 ng of PEAR-GFP plasmid, 1330-1330 ng of pegRNA-mCherry plasmids (one targeting the genome and one targeting the PEAR plasmid), 425-425 ng of sgRNA-mCherry plasmids (one targeting the genome and one targeting the PEAR plasmid) and 2660 ng of PE2 coding plasmid were mixed with 16 μl Turbofect reagent diluted in 800 μl serum-free DMEM. The mixture was then incubated for 30 minutes before it was added to the cells. Three days after transfection cells were trypsinised and sorted directly into genomic lysis buffer, which was followed by genomic DNA extraction.

In all experiments the pCMV-PE2 ^[10]^ plasmid was used as the prime editor expressing plasmid and pX330-Flag-wtSpCas9-H840A as the nSpCas9 and pX330-Flag-dSpCas9 as the dSpCas expressing plasmid. When no second nick was introduced a mock sgRNA expressing plasmid (pAT9922-mCherry or pAT9762-BFP) was used.

### Flow cytometry and cell sorting

Flow cytometry analysis was carried out using an Attune NxT Acoustic Focusing Cytometer (Applied Biosystems by Life Technologies). In all experiments, a minimum of 10,000 viable single cells were acquired by gating based on the side and forward light-scatter parameters. BFP, GFP, mCherry and mScarlet signals were detected using the 405 (for BFP), 488 (for GFP) and 561 nm (for mCherry and mScarlet) diode laser for excitation, and the 440/50 (BFP), 530/30 (GFP), 620/15 (mCherry) and 585/16 nm (mScarlet) filter for emission. Attune Cytometric Software v.2.1.0 was used for data analysis. To compare prime editing in the cell line or on a plasmid Pearson’s correlation coefficient was used.

Cell sorting was carried out on a FACSAria III cell sorter (BD Biosciences). The live single cell fraction was acquired by gating based on the side and forward light-scatter parameters. GFP and mCherry signals were detected using the 488 and 561 nm diode laser for excitation and the 530/30 and 610/20 nm filter for emission, respectively. A minimum of 50,000 cells were sorted in all experiments.

### Genomic DNA purification, genomic PCR and EditR analysis

Genomic DNA from FACS or other experiments was extracted according to the Puregene DNA Purification protocol (Gentra Systems Inc.). The purified genomic DNA was subjected to PCR analysis, conducted with Q5 polymerase and locus specific primers (see **Supplementary Table S2** in Supporting Information). PCR products were gel purified via NucleoSpin Gel and PCR Clean-up kit (Macherey-Nagel) and were Sanger sequenced. Editing efficiencies were quantified by EditR 1.0.9 (https://moriaritylab.shinyapps.io/editr_v10/ ^[39]^.

## Acknowledgements

We thank Dr. György Várady and the FACS core facility for their help with cell sorting. We thank Vanessza Laura Végi for proofreading the manuscript and for the helpful comments on it. We thank Vivien Karl, Ildikó Szűcsné Pulinka, Judit Szűcs, Lilla Burkus and Judit Kálmán for their excellent technical assistance.

## Author contributions

A.T. and E.W. conceived and designed the experiments, interpreted the results and wrote the manuscript with input from all authors. D.A.S performed experiments and interpreted the results. A.T. contributed to molecular cloning and performed experiments on mammalian cells. P. I. K. contributed to molecular cloning and created the PEAR cell lines. E.W. supervised the research. All authors read and approved the final manuscript.

## Supporting information

**Figure S1 –.**
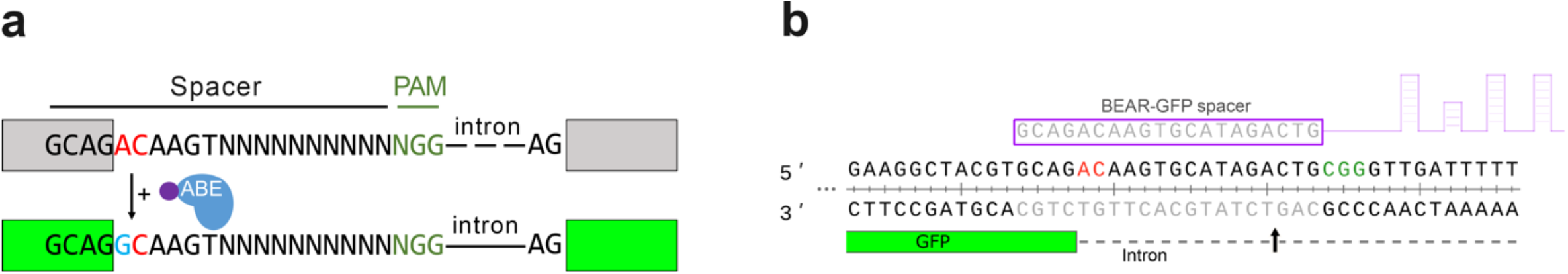
Principle of the Base Editor Activity Reporter (BEAR) assay. (**a**) Schematic representation of the Base Editor Activity Reporter (BEAR). BEAR consists of a split GFP coding sequence separated by an intron, of which the 5’ splice donor site (G-**GT-**AAGT) is altered, resulting in an inactive splice site (dashed line) and a dysfunctional protein (grey). ABE converts the inactive splice site into a functional one. Here the ‘G-**AC-**AAGT’ inactive splice site is illustrated, that can be modified by ABE to ‘G-**GC-**AAGT’, which is a functional non-canonic splice site, and hence, restores GFP expression (green). In this assay, base editors act on the sense strand of the DNA. The altered bases of the splice site are shown in red, the edited base is shown in blue and the variable bases in the sequence of the spacer are shown as ‘N’-s. The PAM sequence is dark green, nCas9 is blue and the fused tadA deaminase is purple. (**b**) Detailed view of the 5’ splice site and the surrounding sequences of the BEAR-GFP plasmid. The 3’ end of the first exon of GFP is shown in green, the intron is shown as a dashed line. The spacer sequence and the target sequence are shown in grey, the PAM is green and the inactive splice site is red. The Cas9 nick site is indicated by a black arrow.

**Figure S2 –.**
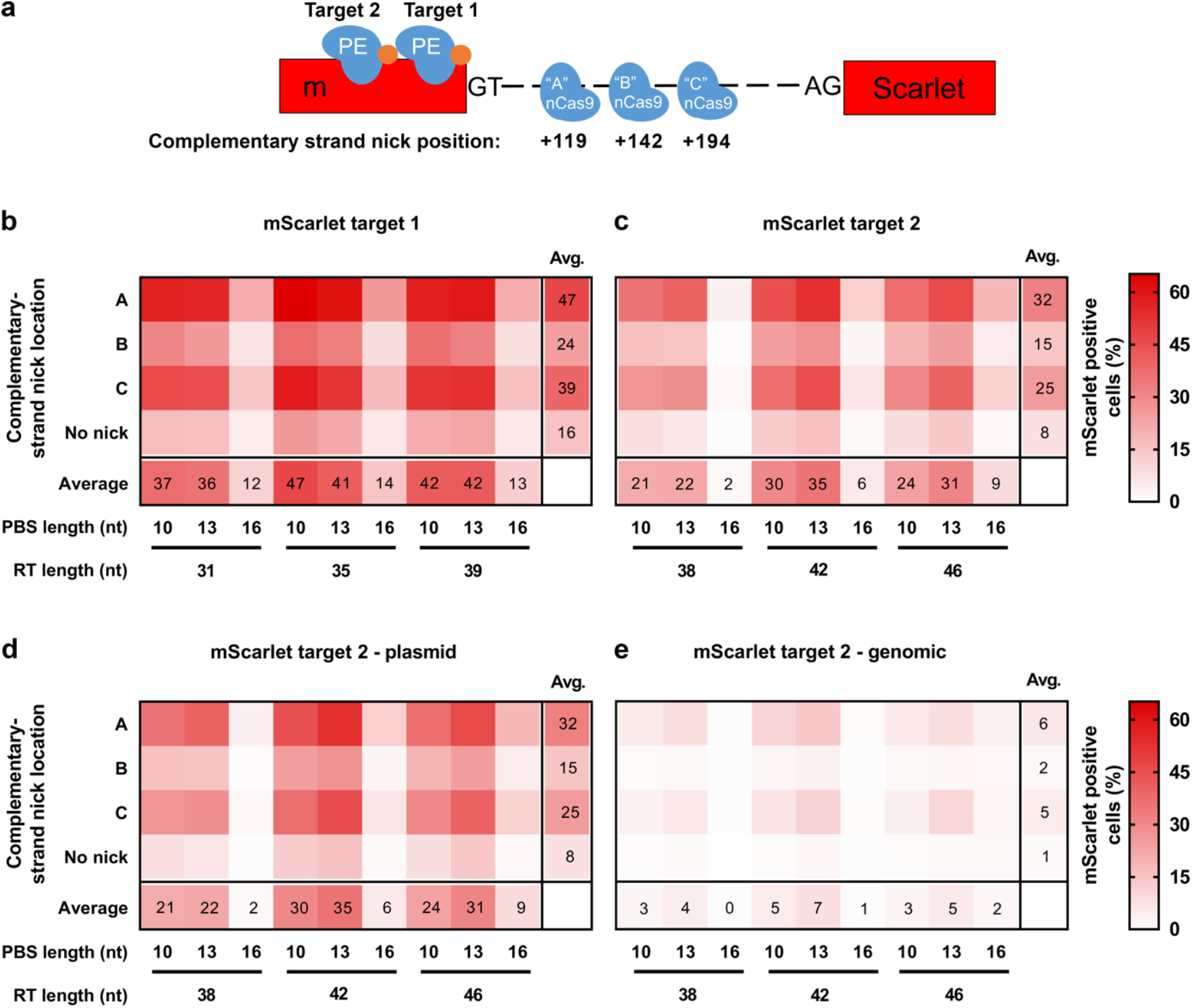
Prime editing of the PEAR-mScarlet plasmid and cell line with two targets. (**a**) Schematic diagram of the PEAR-mScarlet plasmid. The prime editor corrects the inactive splice site and restores mScarlet fluorescence. The second nick positions that enhance prime editing are the same in (**b-e**) and indicated in the figures as A, B and C. The nick positions are given relative to the first nick of target 1, positive values indicate 3’ direction on the targeted DNA. The spacer sequence is directed to either target1 (**b**) or target2 (**c-e**). (**b-e**) The heatmaps show the average percentage of mScarlet positive cells derived from three replicates of the co-transfections of PE with different pegRNAs and nicking sgRNAs when targeting either the PEAR-mScarlet plasmid (**b-d**) or the PEAR-mScarlet cell line (**e**). The spacer sequence is directed to either target 1 (**b**) or target 2 (**c-e**). The position of the second nick is indicated on the left side of the heatmaps with A, B and C letters. When no second nick was introduced, it is indicated as “no nick”.

**Table S1 –.** List of plasmids used in this study.

| Plasmid name | Reference | Comment |
| --- | --- | --- |
| **General** |  |  |
| pCMV-PE2 | Anzalone et al. | Addgene #132775 |
| pX330-Flag-wtSpCas9-H840A | Unpublished | Welker Lab, Addgene #80453 |
| pX330-Flag-dSpCas9 | Kulcsar et al. | Addgene #92113 |
| **PEAR target plasmids** |  |  |
| pDAS12124_PEAR-GFP-1-preedited | This study |  |
| pDAS12125_PEAR-GFP-1 | This study |  |
| PEAR-GFP-2 | Tálas et al. | pAT9651-BEAR-GFP (Addgene #162989) |
| PEAR-mScarlet | Tálas et al. | pAT9752-BEAR-mScarlet (Addgene #162991) |
| pAT9624-BEAR-cloning | Tálas et al. | Addgene #162986 |
| **pegRNA plasmids** |  |  |
| 12067-U6-pegRNA-mCherry | This study | target cloning plasmid |
| 12068-U6-pegRNA-BFP | This study | target cloning plasmid |
| 12227_U6-pegRNA-PEAR-GFP_10PBS-16RT_mCherry | This study | targeting the PEAR-GFP plasmid |
| 12228_U6-pegRNA-PEAR-GFP_13PBS-16RT_mCherry | This study | targeting the PEAR-GFP plasmid |
| 12229_U6-pegRNA-PEAR-GFP_16PBS-16RT_mCherry | This study | targeting the PEAR-GFP plasmid |
| 12230_U6-pegRNA-PEAR-GFP_10PBS-24RT_mCherry | This study | targeting the PEAR-GFP plasmid |
| 12231_U6-pegRNA-PEAR-GFP_13PBS-24RT_mCherry | This study | targeting the PEAR-GFP plasmid |
| 12232_U6-pegRNA-PEAR-GFP_16PBS-24RT_mCherry | This study | targeting the PEAR-GFP plasmid |
| 12233_U6-pegRNA-PEAR-GFP_10PBS-33RT_mCherry | This study | targeting the PEAR-GFP plasmid |
| 12234_U6-pegRNA-PEAR-GFP_13PBS-33RT_mCherry | This study | targeting the PEAR-GFP plasmid |
| 12235_U6-pegRNA-PEAR-GFP_16PBS-33RT_mCherry | This study | targeting the PEAR-GFP plasmid |
| 12248_U6-pegRNA-PEAR-GFP-2_10PBS-22RT_mCherry | This study | targeting the PEAR-GFP-2 plasmid and the PEAR-GFP cell line |
| 12249_U6-pegRNA-PEAR-GFP-2_13PBS-22RT_mCherry | This study | targeting the PEAR-GFP-2 plasmid and the PEAR-GFP cell line |
| 12250_U6-pegRNA-PEAR-GFP-2_16PBS-22RT_mCherry | This study | targeting the PEAR-GFP-2 plasmid and the PEAR-GFP cell line |
| 12251_U6-pegRNA-PEAR-GFP-2_10PBS-26RT_mCherry | This study | targeting the PEAR-GFP-2 plasmid and the PEAR-GFP cell line |
| 12252_U6-pegRNA-PEAR-GFP-2_13PBS-26RT_mCherry | This study | targeting the PEAR-GFP-2 plasmid and the PEAR-GFP cell line |
| 12253_U6-pegRNA-PEAR-GFP-2_16PBS-26RT_mCherry | This study | targeting the PEAR-GFP-2 plasmid and the PEAR-GFP cell line |
| 12254_U6-pegRNA-PEAR-GFP-2_10PBS-33RT_mCherry | This study | targeting the PEAR-GFP-2 plasmid and the PEAR-GFP cell line |
| 12255_U6-pegRNA-PEAR-GFP-2_13PBS-33RT_mCherry | This study | targeting the PEAR-GFP-2 plasmid and the PEAR-GFP cell line |
| 12256_U6-pegRNA-PEAR-GFP-2_16PBS-33RT_mCherry | This study | targeting the PEAR-GFP-2 plasmid and the PEAR-GFP cell line |
| 12257_U6-pegRNA-PEAR-mScarlet-target1_10PBS-31RT_BFP | This study | targeting the PEAR-mScarlet - target1 plasmid and the PEAR-mScarlet cell line |
| 12258_U6-pegRNA-PEAR-mScarlet-target1_13PBS-31RT_BFP | This study | targeting the PEAR-mScarlet - target1 plasmid and the PEAR-mScarlet cell line |
| 12259_U6-pegRNA-PEAR-mScarlet-target1_16PBS-31RT_BFP | This study | targeting the PEAR-mScarlet - target1 plasmid and the PEAR-mScarlet cell line |
| 12260_U6-pegRNA-PEAR-mScarlet-target1_10PBS-35RT_BFP | This study | targeting the PEAR-mScarlet - target1 plasmid and the PEAR-mScarlet cell line |
| 12261_U6-pegRNA-PEAR-mScarlet-target1_13PBS-35RT_BFP | This study | targeting the PEAR-mScarlet - target1 plasmid and the PEAR-mScarlet cell line |
| 12262_U6-pegRNA-PEAR-mScarlet-target1_16PBS-35RT_BFP | This study | targeting the PEAR-mScarlet - target1 plasmid and the PEAR-mScarlet cell line |
| 12263_U6-pegRNA-PEAR-mScarlet-target1_10PBS-39RT_BFP | This study | targeting the PEAR-mScarlet - target1 plasmid and the PEAR-mScarlet cell line |
| 12264_U6-pegRNA-PEAR-mScarlet-target1_13PBS-39RT_BFP | This study | targeting the PEAR-mScarlet - target1 plasmid and the PEAR-mScarlet cell line |
| 12265_U6-pegRNA-PEAR-mScarlet-target1_16PBS-39RT_BFP | This study | targeting the PEAR-mScarlet - target1 plasmid and the PEAR-mScarlet cell line |
| 12266_U6-pegRNA-PEAR-mScarlet-target2_10PBS-38RT_BFP | This study | targeting the PEAR-mScarlet - target2 |
| 12267_U6-pegRNA-PEAR-mScarlet-target2_13PBS-38RT_BFP | This study | targeting the PEAR-mScarlet - target2 |
| 12268_U6-pegRNA-PEAR-mScarlet-target2_16PBS-38RT_BFP | This study | targeting the PEAR-mScarlet - target2 |
| 12269_U6-pegRNA-PEAR-mScarlet-target2_10PBS-42RT_BFP | This study | targeting the PEAR-mScarlet - target2 |
| 12270_U6-pegRNA-PEAR-mScarlet-target2_13PBS-42RT_BFP | This study | targeting the PEAR-mScarlet - target2 |
| 12271_U6-pegRNA-PEAR-mScarlet-target2_16PBS-42RT_BFP | This study | targeting the PEAR-mScarlet - target2 |
| 12272_U6-pegRNA-PEAR-mScarlet-target2_10PBS-46RT_BFP | This study | targeting the PEAR-mScarlet - target2 |
| 12273_U6-pegRNA-PEAR-mScarlet-target2_13PBS-46RT_BFP | This study | targeting the PEAR-mScarlet - target2 |
| 12274_U6-pegRNA-PEAR-mScarlet-target2_16PBS-46RT_BFP | This study | targeting the PEAR-mScarlet - target2 |
| 12239_U6-pegRNA-RNF2_mCherry | This study | Targeting RNF2 genomic target |
| 12241_U6-pegRNA-FANCF_mCherry | This study | Targeting FANCF genomic target |
| 12244_U6-pegRNA-HEK3_mCherry | This study | Targeting HEK3 genomic target |
| **sgRNA plasmids** |  |  |
| 12136_U6-sgRNA-PEAR-GFP_nick(+89)_mCherry | This study | Secondary nick on the PEAR-GFP plasmid |
| 12137_U6-sgRNA-PEAR-GFP_nick(+17)_mCherry | This study | Secondary nick on the PEAR-GFP plasmid |
| 12138_U6-sgRNA-PEAR-GFP_nick(-59)_mCherry | This study | Secondary nick on the PEAR-GFP plasmid |
| 12139_U6-sgRNA-PEAR-GFP_nick(-100)_mCherry | This study | Secondary nick on the PEAR-GFP plasmid |
| 12140_U6-sgRNA-PEAR-GFP_nick(-162)_mCherry | This study | Secondary nick on the PEAR-GFP plasmid |
| 12198_U6-sgRNA-PEAR-GFP-2_nick(+103)_mCherry | This study | Secondary nick on the PEAR-GFP-2 plasmid, and in the PEAR-GFP cell line |
| 12199_U6-sgRNA-PEAR-GFP-2_nick(+126)_mCherry | This study | Secondary nick on the PEAR-GFP-2 plasmid, and in the PEAR-GFP cell line |
| 12200_U6-sgRNA-PEAR-GFP-2_nick(+17)_mCherry | This study | Secondary nick on the PEAR-GFP-2 plasmid, and in the PEAR-GFP cell line |
| 12210_U6-sgRNA-PEAR-mScarlet_nick(+103)_BFP | This study | Secondary nick on the PEAR-mScarlet plasmid and in the PEAR-mScarlet cell line |
| 12211_U6-sgRNA-PEAR-mScarlet_nick(+126)_BFP | This study | Secondary nick on the PEAR-mScarlet plasmid and in the PEAR-mScarlet cell line |
| 12212_U6-sgRNA-PEAR-mScarlet_nick(+17)_BFP | This study | Secondary nick on the PEAR-mScarlet plasmid and in the PEAR-mScarlet cell line |
| 12156_U6-sgRNA-RNF2_nick(+41)_mCherry | This study | Secondary nick in the genome |
| 12177_U6-sgRNA-FANCF_nick(+48)_mCherry | This study | Secondary nick in the genome |
| 12178_U6-sgRNA-HEK3_nick(+90)_mCherry | This study | Secondary nick in the genome |
| pAT9922_U6-sgRNA-mock-mCherry | This study | mock sgRNA used in when not nicking the plasmid/genome |
| 9762_U6-sgRNA-mock-TagBFP | This study | mock sgRNA used in when not nicking the plasmid/genome |

**Table S2 –.**
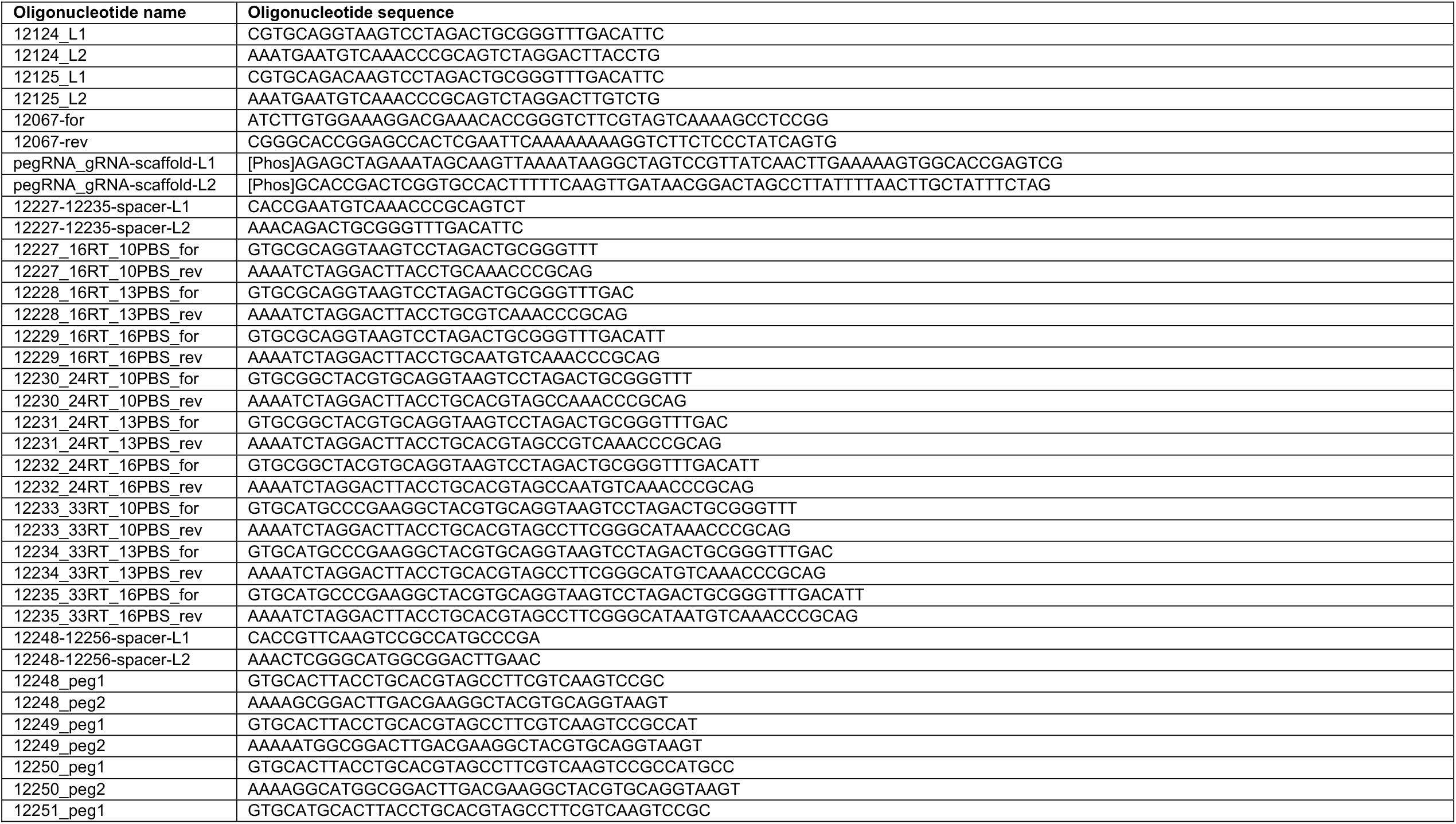
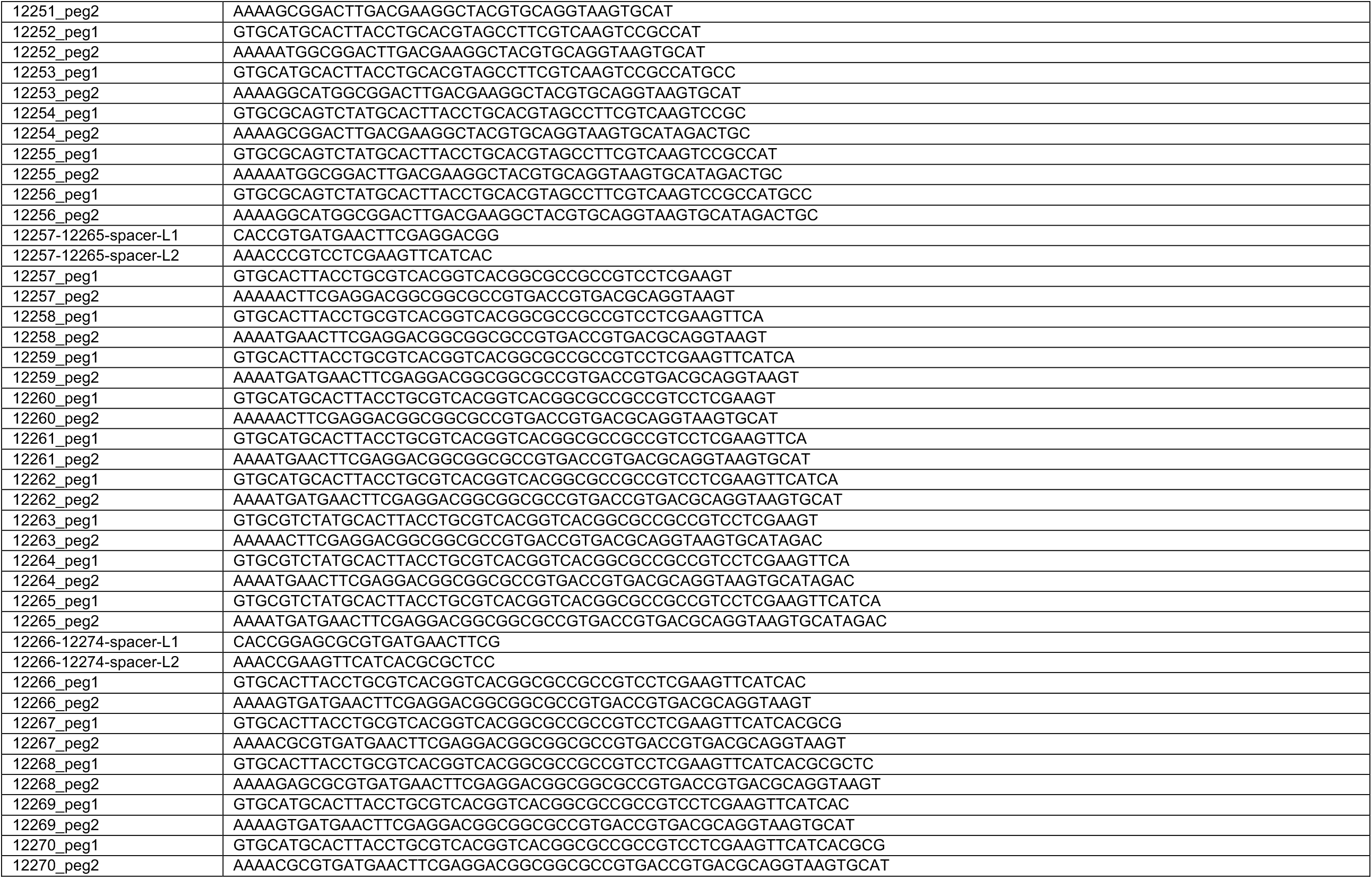
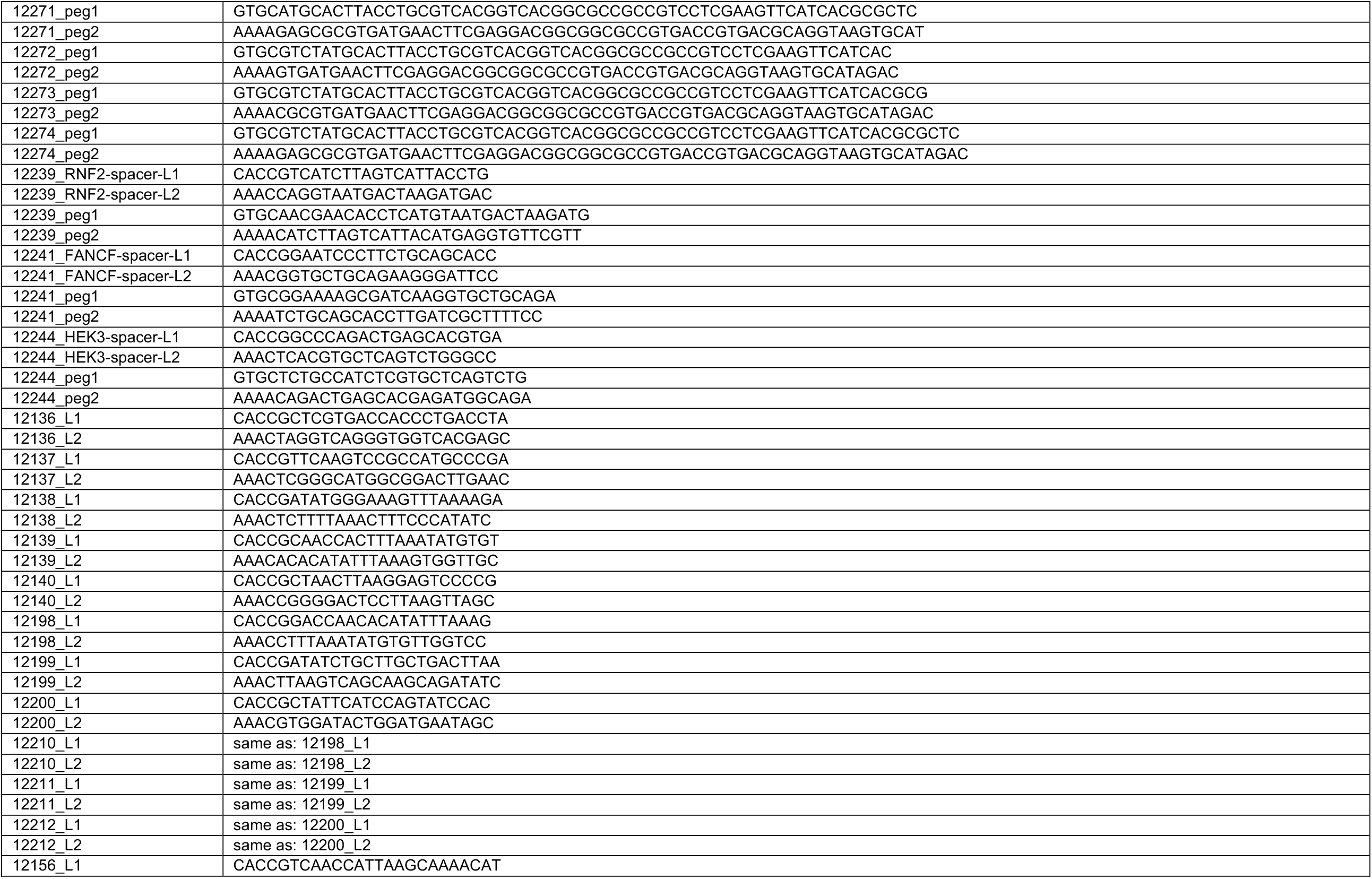
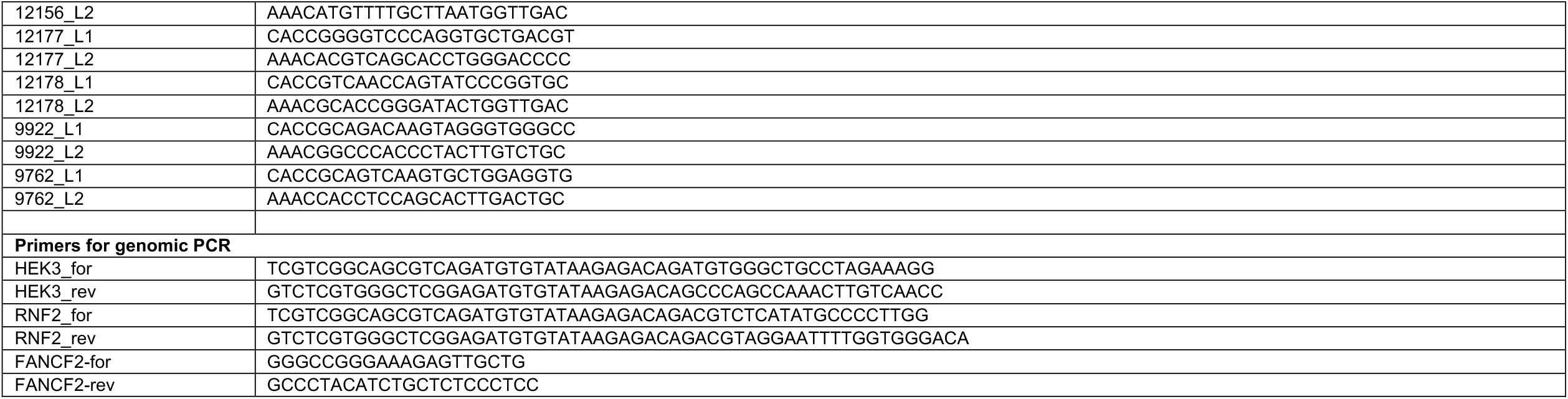
List of oligonucleotides used in this study.

